# Long days induce adaptive secondary dormancy in seed of the Mediterranean plant *Aethionema arabicum*

**DOI:** 10.1101/2024.01.08.574645

**Authors:** Zsuzsanna Mérai, Kai Graeber, Fei Xu, Mattia Dona, Katarina Lalatović, Per K.I. Wilhelmsson, Noe Fernandez-Pozo, Stefan A. Rensing, Gerhard Leubner-Metzger, Ortrun Mittelsten Scheid, Liam Dolan

## Abstract

Secondary dormancy is an adaptive trait that increases reproductive success by aligning seed germination with permissive conditions for seedling establishment. *Aethionema arabicum* is an annual plant and member of the Brassicaceae that grows in environments characterized by hot and dry summers. *Aethionema arabicum* seeds may germinate in early spring when seedling establishment is permissible. We demonstrate that long-day light regimes induce secondary dormancy in seed of *Aethionema arabicum* (CYP accession) repressing germination in summer when seedling establishment is riskier. Characterization of mutants screened for defective secondary dormancy demonstrated that RGL2 mediates repression of genes involved in GA signalling. Exposure to high temperature alleviates secondary dormancy, restoring germination potential. These data are consistent with the hypothesis that long-day-induced secondary dormancy and its alleviation by high temperatures, may be part of an adaptive response limiting germination to conditions permissive for seedling establishment in spring and autumn.

## INTRODUCTION

Long-term survival of many plants depends on the alignment of seed germination to seasonally changing conditions. Seed dormancy, defined as a failure of viable seeds to germinate under otherwise suitable conditions, is an important component of this adaptation. Primary dormancy develops during seed maturation and can be influenced by physiological, physical, or morphological factors, like phytohormone balance, hard seed coat, or immature embryos, respectively ^1,2^. Primary dormancy prevents early germination before shedding or schedules germination of shed seeds to the optimal time window ^3^. Secondary seed dormancy can be induced after shedding in non-dormant imbibed seeds by environmental stimuli like suboptimal temperature, hypoxia, or darkness ^4–9^. Unless the secondary dormancy is released by antagonistic signals, seeds remain dormant even in conditions permissive for germination, indicating a memory effect as part of a dormancy cycle.

Despite the importance of controlled and uniform germination timing for maximum crop production, little is known about the mechanism by which secondary dormancy is induced or alleviated during dormancy cycling ^4,10,11^. Dormancy cycling is often the result of the interaction of multiple factors. For example, darkness in combination with far-red light pulses can induce secondary dormancy in *Arabidopsis thaliana*, but nothing else is known about mechanisms promoting or repressing secondary dormancy ^8^.

Here we demonstrate that exposure to light induces secondary dormancy in *Aethionema arabicum* (*Brassicaceae*). Light induction of secondary dormancy results from a decrease in GA signalling and requires the activity of the DELLA protein RGL2. Secondary dormancy is alleviated by aridity and high temperatures. We propose that the induction of dormancy by long days and its alleviation by a combination of drought and high temperatures, may be an adaptive mechanism that increases the probability of seeds germinating in conditions that are permissive for seedling establishment.

## RESULTS

### Light induces secondary seed dormancy in Aethionema arabicum seeds

Since light inhibits germination in non-dormant *Ae. arabicum* seed of the Cyprus accession (CYP) ^12,13^, we hypothesized that light can also induce secondary dormancy. We exposed six-month-old, after-ripened, imbibed CYP seed populations to 24 h of light per day (24 L) for 7 days or 24 h of darkness (24 D) for 7 days (control). While 98% of seed exposed to 24 h darkness for 7 days germinated, almost no seed germinated when exposed to light. 0.7% of the seed population germinated when exposed to 100 µmol m^-2^ s^-1^ light and 0% of seed germinated in 230 µmol m^-2^ s^-1^ light (Fig. 1 A). These data are consistent with the previously published results that light inhibits seed germination in the CYP accession of *Ae. arabicum* (Merai et al 2019, 2023). We refer to this phenomenon as direct light-inhibition of germination (also known as photoinhibition). To specifically test the hypothesis that light induces secondary dormancy – dormancy in seed when grown in permissive germination conditions after receiving a transient dormancy-inducing signal – after-ripened CYP seed were grown under 24 h constant light for periods of between 1 and 7 days before transfer to 24 h darkness for 7 days. If light induced secondary dormancy, we expected that a period of light exposure would inhibit germination in the subsequent dark period, which is permissive for germination. Almost 100% of seed exposed to 1 day of 24 h light before transfer to 24 h darkness for 6 days germinated, indicating that a short period of light exposure did not induce secondary dormancy. However, longer periods of light treatment before transfer to darkness resulted in lower germination rates. For example, 4.3% and 2.4% of seed germinated after treatment with 7 days of 24 h light before transfer to 24 h dark for a further 7 days in two different light intensities. The inhibition of germination in after-ripened seed by exposure to light lasting after return to darkness conditions permissive for germination, demonstrates that the light treatment induced secondary dormancy in the CYP accession of *Ae. arabicum* (Fig. 1 A).

**Figure 1.**
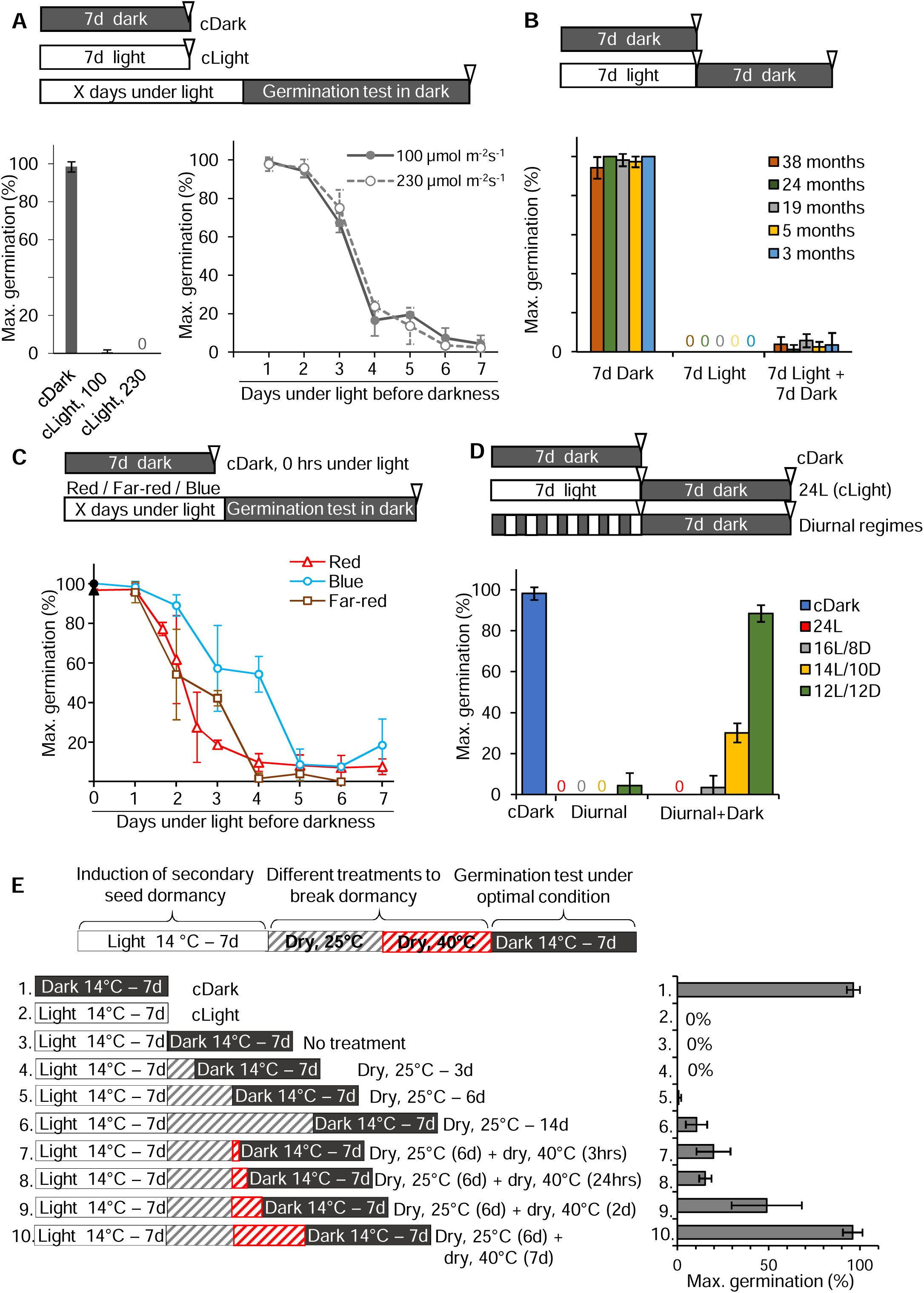
Light induces secondary seed dormancy in *Ae. arabicum*. **(A)** Seeds tested for maximal germination. Top: scheme of light regimes. Left: germination under constant darkness (cDark), constant white light at 100 µmol m^-2^s^-1^ (cLight, 100), or 230 µmol m^-2^s^-1^ (cLight, 230) intensities for 7 days. Right: seeds kept under constant light for 1-7 days before plates were transferred to darkness. Maximal germination was scored after 7 days in darkness. **(B)** Top: scheme of light regimes. Below: five seed batches at different times after harvest were tested for maximal germination after 7 days in darkness, after 7 days in constant light, or after 7 days in constant light followed by 7 days of darkness. **(C)** Maximal seed germination after keeping the seeds under red, blue, or far-red light for 0-7 days, followed 7 days in darkness. 0 time point indicates that seeds were kept under constant darkness for 7 days as a control. **(D)** Seeds were kept in darkness (cDark), or constant light (24L), or diurnal regimes for 7 days as schematically indicated. Maximal seed germination was scored after transferring the plates to darkness for additional 7 days. **(E)** Left: scheme of the treatments to test the alleviation of secondary dormancy induced by 7 days exposure to constant light as described in C. Right: after each treatment, maximal germination capacity of rehydrated seeds was determined after 7 days in darkness. Error bars indicate the standard deviation of three biological replicates.

We tested if light-induced secondary dormancy depends on the length of the after-ripening period (seed age). After-ripening usually occurs within three months of harvesting and seed that are less than three months old display primary dormancy (Fig. S1A) (Merai et al 2019, 2023). We tested the effect of light exposure on seed that had been stored 3 to 38 months after harvest. We first showed that 94-100% of five different seed batches germinated equally well in permissive conditions (constant darkness), which verified that they were not primary dormant (Fig. 1 B). No seed germinated under 24 h light for 7 days and few (max. 5.8%) germinated after being exposed to 24 h light for 7 days before transfer to darkness (Fig. 1 B). Taken together, these data indicate that light-induced secondary dormancy in after-ripened seed is independent of seed age.

Since white light induced secondary dormancy, we set out to identify the wavelengths of light that induce secondary dormancy. Post ripened seed were sown and grown under different light regimes; red light (654 nm peak), far-red light (735 nm peak), and blue light (441 nm peak). Secondary dormancy was induced by red, far-red, or blue light (Fig. 1 C). However, blue light was less efficient than the other wavelengths (Fig. 1 C). These data suggests that any of the light wavelengths sensed by the red or blue light photoreceptors induce secondary dormancy.

Having shown that exposure to 24 h light per day (24L) for 7 days induced secondary dormancy, we tested if the relative length of the light and dark periods in a 24-hour diurnal cycle impacted the onset of secondary dormancy. We exposed seeds to long day – 16 h light (16L) / 8 h dark (8D), 14L / 10D – or 12L / 12D diurnal cycles. Secondary dormancy was induced in 24L (continuous light) and 16L/8D conditions where 100% and 97.6% of the seed did not germinate, respectively (Fig. 1 D). 30.2% of seed did not germinate in the 14L/10D diurnal cycle. However, there was almost no secondary dormancy in 12L/12D conditions where 88% seed germinated in the dark period that followed the light exposure (Fig. 1 D). These data indicate that secondary dormancy is induced by long-day, diurnal light-dark regimes.

As long day conditions induced secondary dormancy, we asked how this might be alleviated once favorable conditions for germination return. Given the environment in which *Ae. arabicum* grows in Cyprus – hot, dry summers – we hypothesized that high temperature and aridity would alleviate secondary dormancy. To test this hypothesis, seed with secondary dormancy induced by 7 days of continuous light (24L) during imbibition were afterwards exposed to aridity and high temperature. For this, seeds were placed on dry filter paper without water and exposed to different periods at 25°C or 40°C in the dark (Fig. 1 E). After exposure to these different temperature regimes, the germination capacity of re-wetted seeds was tested in permissive conditions (darkness) at 14°C, the optimal germination temperature (Fig. 1 E). While the 25°C treatment partially restored germination (10.3%), the 40°C treatment restored full germination (95.8%). These data indicate that high temperature alleviates light-induced secondary dormancy. Taken together, these data demonstrate that long-day conditions induce secondary dormancy in *Ae. arabicum*, and that this dormancy can be alleviated by exposure to high temperature.

### RGL2 is required for the establishment of secondary dormancy in Ae. arabicum

To identify genes required for the induction of secondary dormancy, we screened a gamma radiation-mutated seed population of *Ae. arabicum* CYP accession for mutants that germinate despite being exposed to light conditions that induce secondary dormancy in wild type. A minimum of 20 imbibed seeds per M2 or M3 family was plated on wet filter paper and illuminated with white light for 24 h for 7 days to induce secondary dormancy (Fig. 2 A). Seeds were then transferred to 24 h darkness for a further 7 days. In contrast to non-germinating wild type, almost 100% of seeds from the p24H4 M3 family germinated in the following darkness period (Fig. 2 A,B). The mutant was backcrossed to a CYP wild-type plant and the F_1_ plants self-fertilized to generate an F_2_ population. Approximately 25% of the F_2_ seeds germinated after induction of secondary dormancy (defective dormancy phenotype), whereas approximately 75% did not germinate under these conditions (wild type dormancy phenotype) (Fig. S3A). This indicates that the absence of secondary dormancy in the mutant line was inherited as a single, recessive Mendelian trait (Fig. 2 A, Fig. S2A, S3A). Whole-genome sequencing of DNA isolated from bulked wild type seedlings and bulked seedlings with the mutant phenotype from a segregating F_2_ population identified a four base pair deletion and a single nucleotide change in *Aa31LG5G13950* in 93.4% of mutant bulk population. The frequency of this allele in the bulked wild type sibling population was 36.7% (expected frequency is 33.3% if the mutation caused a recessive phenotype) (Fig. S2B). The secondary dormancy defect co-segregated with the mutations in *Aa31LG5G13950,* consistent with the hypothesis that mutations in this gene cause the defective dormancy (Fig. S3A). The mutation generates a frameshift at the end of the coding region that predicts the production of a protein that is 75 aa longer than in the wild type (Fig. 2 C, Fig. S2). The predicted protein sequence of *Aear31LG5G13950* is similar to the Arabidopsis RGL2 protein, a member of the DELLA family. A blast search with all five Arabidopsis DELLA proteins identified four DELLA proteins in *Ae. arabicum*. Amino acid sequence alignment and phylogenetic analysis confirmed that *Aear31LG5G13950* encodes the RGL2 ortholog and was therefore designated *AearRGL2* (Fig. S3 B,C). The defective secondary dormancy phenotype of mutants homozygous for the loss-of-function mutation in *AearRGL2* demonstrates that AearRGL2 is required for the initiation of light induced secondary dormancy in *Ae. arabicum*. A previous report showed the role of RGL2 in the dark-induced secondary dormancy in Arabidopsis ^8^. Our data suggest a conserved role of RGL2 in secondary dormancy establishment even though with converse initiation signal; not darkness but light promotes secondary dormancy in *Ae. arabicum*.

**Figure 2.**
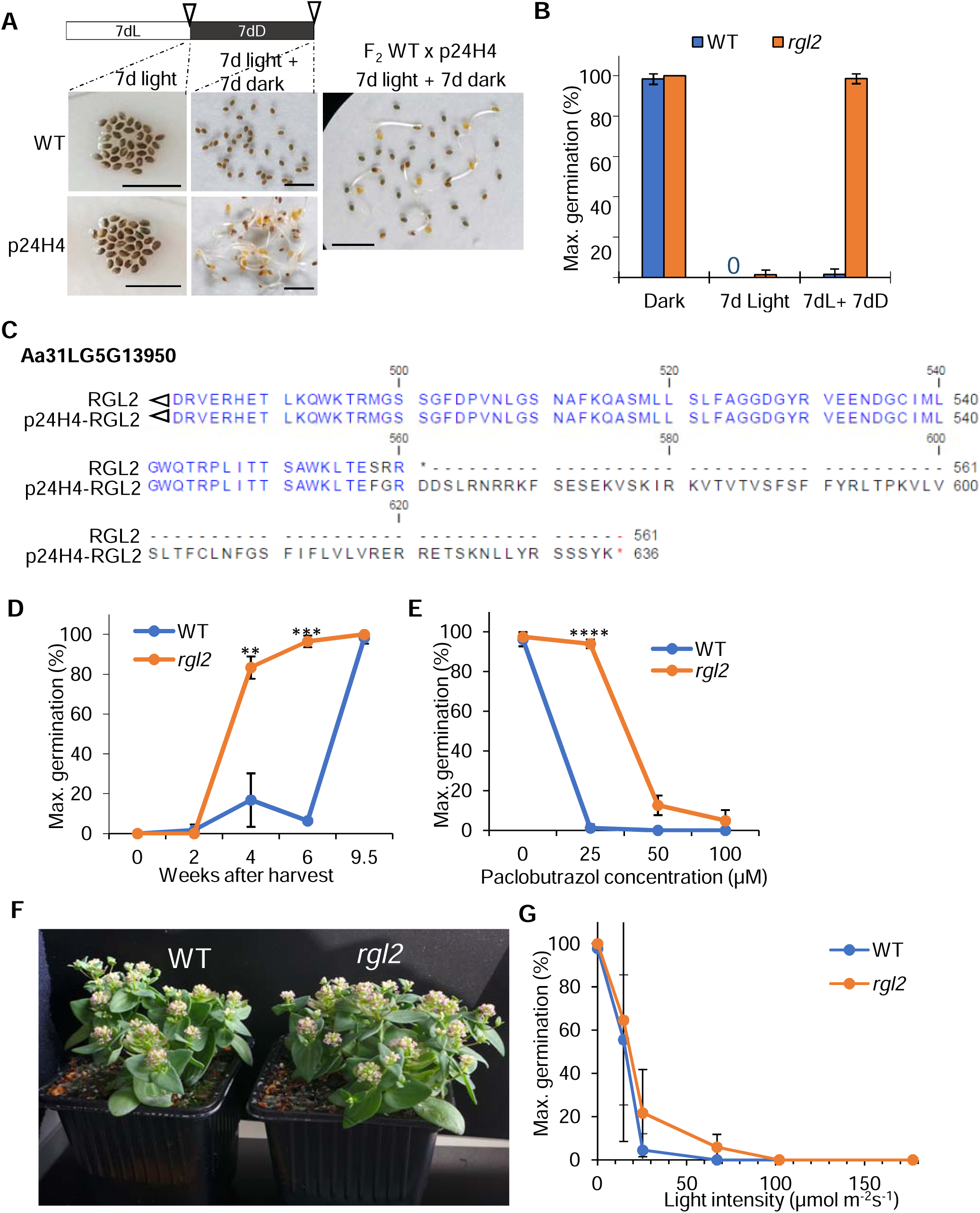
RGL2 mediates the entrance of secondary dormancy. **(A)** Germination phenotype of WT and p24H4 seeds and a segregating F2 population from a cross between them. Scale bars represent 1 cm. **(B)** Maximal germination of WT and p24H4 seeds after exposing them to the light regime in A. **(C)** C-terminal end of the translated protein sequence of RGL2 (*Aa31LG5G13950*) from the WT and p24H4 mutant cDNA. **(D, E)** Maximal germination scored after dark incubation for 7 days. Significant differences are indicated by asterisks, ** *P*< 0.01, *** *P*< 0.001, **** *P*< 0.0001 tested by Welch test in pairwise comparison. **(F)** 9-week-old WT and *rgl2* mutant plants. **(G)** Maximum germination after exposing WT and mutant seeds to different white light intensities for 7 days. Error bars: standard deviation of three biological replicates.

In addition to the role of RGL2 in secondary dormancy, there are two phenotypes described for the *rgl2* single mutant in Arabidopsis; weaker primary dormancy in seeds and higher tolerance of paclobutrazol (PAC), an inhibitor of gibberellin (GA) biosynthesis, during seed germination ^8,14^. The *Ae. arabicum rgl2* mutant seed phenotypes are similar to the Arabidopsis *rgl2* mutant phenotypes; *Ae. arabicum rgl2* mutant seeds release primary dormancy sooner than wild type, and mutants are resistant to PAC compared to wild type (Fig. 2 D, E). Like Arabidopsis *rgl2* mutants, *Ae. arabicum rgl2* plants are morphologically indistinguishable from wild type (Fig. 2 F). The gemination of the *rgl2* mutant and wild type seed was inhibited by 24 h light (100 µmol m^-2^ s^-1^) for 7 days constant white light (Fig. 2 G), indicating RGL2 is not required for direct light-inhibition of germination. Taken together, these data demonstrate that RGL2 is required for light-induced secondary dormancy in *Ae. arabicum* and contributes to repressing primary dormancy, as in Arabidopsis.

### Transcriptomes identify genes and gene sets associated with secondary seed dormancy

To identify mechanisms that promote secondary dormancy, we set out to identify genes and gene sets that were differentially expressed during light-induced secondary dormancy. We hypothesized that these genes would be differentially represented in transcriptomes generated from wild type seed exposed to 24 h light for one day (light-responding, but not yet dormant) and wild type seeds exposed to 24 h light for 7 days (secondary dormant) (Fig. 3 A). Similarly, we hypothesized that genes associated with light-induced secondary dormancy would be differentially represented in transcriptomes generated from wild type (secondary dormant) and *rgl2* (non-dormant) seed exposed to 24 h light for 7 days (Fig. 3 A). Identifying genes that were differentially expressed in both comparisons would define a suite of differentially expressed genes characteristic of secondary dormancy. The comparison identified a total of 3,330 differentially expressed genes (DEGs) associated with secondary dormancy.

**Figure 3.**
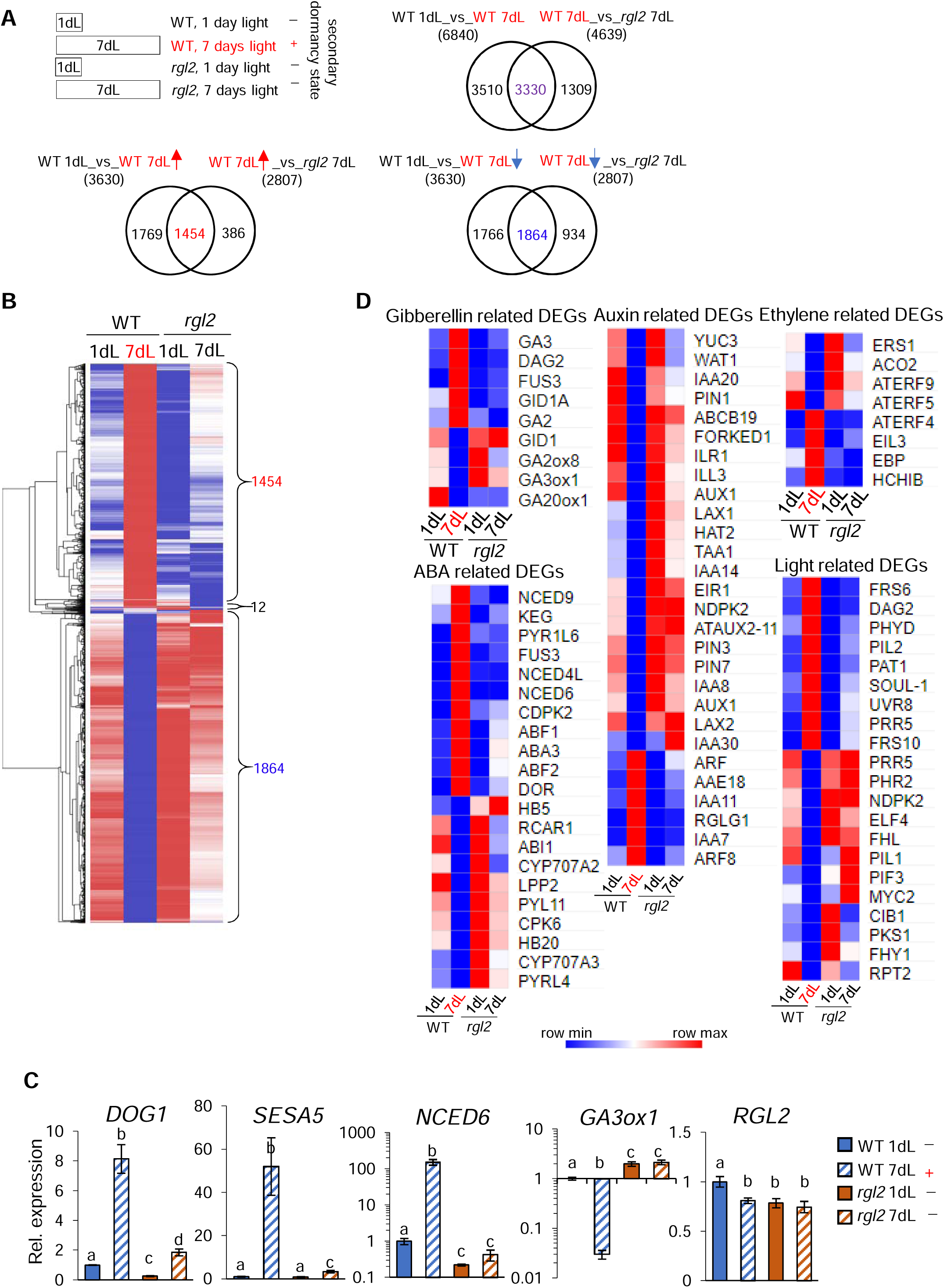
Different transcriptional responses at the light-induced secondary seed dormancy in wild type versus *rgl2* mutant. **(A)** Experimental design. Red letters: samples with established light-induced secondary dormancy. Venn diagrams show the number of differentially regulated genes (DEGs) in the respective comparison. **(B)** Heatmap indicating the relative expression of all 3330 common DEGs. Coloring is based on the log2 transformation of reads per kilobase of transcript per million mapped reads (RPKM) values by row. Shades of blue indicate lower expression and shades of red indicate higher expression. **(C)** Quantitative RT-PCR results for selected genes. The expression level of WT 7dL, *rgl2* 1dL, and *rgl2* 7dL is presented as fold change relative to the average expression level of WT 1dL set to 1. Letters indicate significant differences with \**P*<0.05 (Welch test). Error bars: SD of 3 biological replicates. **(D)** Heatmap indicating relative expression of selected DEGs related to hormonal or light regulation. Colors as in B.

To validate that these 3,330 genes included genes associated with seed dormancy, we screened the list for genes whose orthologs function in seed dormancy and germination in Arabidopsis. This search identified genes including *AearDOG1* (*DELAY OF GERMINATION 1),* two *NINE-CIS-EPOXYCAROTENOID DIOXYGENASE* (*NCED*) genes (*AearNCED6,* and *AearNCED9*) and one *NINE-CIS-EPOXYCAROTENOID DIOXYGENASE-LIKE* (*NCEDL*) (*AearNCED4L)* involved in abscisic acid (ABA) synthesis, the *SEED STORAGE ALBUMIN 5* (*AearSESA5*), and *GIBBERELLIN 3-OXIDASE 1* (*AearGA3ox1*) involved in gibberellin (GA) biosynthesis ^15–18^. The differential expression of many of these genes between secondary dormant and non-secondary dormant seed was verified by reverse transcriptase quantitative polymerase chain reaction (RT-qPCR) (Fig. 3 C, Table S1). Therefore, we conclude that this suite of 3,330 differentially expressed genes identifies genes associated with the seed dormancy and are therefore referred to them as dormancy-associated genes (Fig. 3 A-C, Table S1).

Positive regulators of dormancy would accumulate at higher levels only at day 7 in the wild type but not in the other three compared samples. This is the case for 1,454 mRNAs. By contrast, the levels of mRNA for negative regulators of dormancy would be expected to be the lowest at day 7 in the wild type and be higher in the other three samples. 1,864 mRNAs fall in this category (Fig. 3 A-C; Table S1).

Approximately one third (1,115) of the dormancy-associated genes (3,330) were also differentially expressed between *rgl2* mutant seed treated with 24 h light for 1 day and *rgl2* mutants seed treated with 24 h light for 7 days (Fig. S4 A,C). The corresponding mRNAs also accumulated at lower levels at day 7 in the *rgl2* mutant than in wild type (Fig. S4). This indicates that RGL2 represses the accumulation of these genes during the onset of secondary dormancy. This is exemplified by *AearDOG1* and *AearSESA5.* The expression of these two genes in the RNAseq experiment increased 5.3 and 118-fold between 1 and 7 days of light in wild type, respectively (Fig. S4 B). However, the change in expression between 1 and 7 days of light was only 3.6 and 5.8-fold, respectively, in the *rgl2* mutant (Fig. 3 C, Fig. S4 B). Furthermore, *RGL2* was not among this group of 3,330 differentially expressed genes. Steady state levels of RGL2 mRNA were identical in wild type and *rgl2* at 7 day light treatment. This indicates that, while *RGL2* controls the expression of genes associated with dormancy, it is not induced by the dormancy-inducing light treatment, and it does not regulate its own expression (Fig. 3 C).

The transcriptome differences between dormant and non-dormant seed support the hypothesis that RGL2 regulates gene expression during the onset of secondary dormancy. Gibberellins (GA) promote germination and inhibit dormancy in seeds of many species and gibberellin 3-oxidase 1 is an enzyme involved in GA synthesis. Among the differentially expressed genes, *AearGA3ox1* mRNA levels were significantly higher in non-dormant than dormant seeds; steady state levels of *AearGA3ox1* mRNA in *rgl2* were twice the level in wild type at day 1, consistent with the known repressive function of RGL2 on *GA3ox1* gene expression. Importantly, the expression was 70 times lower at day 7 than at day 1 in wild type (Fig. 3 C). This indicates that RGL2 represses the expression of this gene during the onset of secondary dormancy. By contrast, ABA inhibits germination and promotes dormancy in seeds of many species. Steady state levels of mRNAs encoding two ABA catabolic enzymes –AearCYP707A2 and AearCYP707A3 – and a repressor of ABA signalling – *AearABI1* – were higher in *rgl2* mutant than in wild type at both day 1 and day 7 of light treatment (Fig. 3 D). This indicates that RGL2 represses genes encoding enzymes that reduce ABA levels and repressors ABA signalling (AearCYP707A2, AearCYP707A3, and AearABI1) (Fig. 3 D). While these modulators of ABA activity are regulated by RGL2, other previously identified regulators are not. For example, the mRNA levels of ABA signalling regulators, *AearABI3* or *AearABI5* ^17,19,20^, were lower after dormancy induction in wild type and expressed at even lower levels in the *rgl2* mutant (Fig. S4 D). This suggests that light-induced dormancy is independent of *AearABI3* or *AearABI5* function, at least at the level of mRNA accumulation (Fig. S4 D). The higher steady state levels of mRNAs encoding promoters of GA biosynthesis and mRNAs encoding repressors of ABA function in *rgl2* mutants are consistent with the hypothesis that RGL2-regulated secondary dormancy is promoted by ABA and repressed by GA.

Seed dormancy is positively regulated by auxin and mutants with reduced auxin biosynthesis develop weaker primary dormancy in Arabidopsis ^21^. To determine if auxin impacted secondary dormancy, changes in the steady state levels of mRNAs encoding proteins that function in auxin synthesis and transport were investigated. Steady state levels of mRNAs encoding proteins with auxin biosynthesis and transport activities – including *AearTAA1*, *AearYUC3*, *AearPIN1*, *AearPIN3*, *AearPIN7*, *AearAUX1*, *AearLAX1*, *AearLAX2* – were lower in the 7-day light-treated wild type seed (dormant) than at day 1 (non-dormant). This suggests that expression of these genes is repressed when secondary dormancy is induced. Furthermore, steady state levels of these mRNAs were lower in wild type (dormant) than in *rgl2* mutants (non-dormant) seed exposed to 24 h light for 7 days (Fig. 3 D). Taken together, these data indicate that RGL2 represses the expression of genes involved in auxin biosynthesis and transport during the onset of secondary dormancy.

While the analysis of transcriptomes demonstrated that secondary dormancy is associated with repression of genes involved in GA synthesis and genes that reduce ABA levels, we identified other processes associated with secondary dormancy. For an unbiased analysis, we applied gene set enrichment analysis (GSEA) that relies on the fold changes in pair-wise comparisons of all expressed genes. Each gene set includes genes that function in similar biological processes based on gene ontology. We identified 29 common gene sets in the two comparisons (Fig. S5, S6, Table S2, Table S3). Steady state levels of mRNA for two lipid storage and terpene metabolic gene sets (including *AearNCED6, AearNCED9* and *AearNCED4L*) were more abundant in the wild type (dormant) than in non-dormant samples (Fig. S5, S6, Table S2, Table S3). Reciprocally, these two gene sets were expressed at lower levels in *rgl2* mutants (non-dormant) than in the wild type (dormant) seed exposed to 24 h light for 7 days (Fig. S5, S6, Table S2, Table S3). Among the 27 gene sets whose mRNAs were less abundant in secondary dormant than in non-dormant seed were ribosomal biogenesis-related processes, translation, DNA replication, and RNA metabolic processing (Fig. S5, S6 Table S2, Table S3). This indicates that sets of genes encoding proteins active in general metabolic activity and growth are repressed during the onset of secondary dormancy.

Taken together, the data from transcriptomes demonstrate that RGL2 promotes the expression of genes encoding ABA biosynthesis and represses genes encoding GA biosynthesis activities during the onset of secondary dormancy. The roles played by these hormones during secondary dormancy is similar to their role in primary dormancy where ABA promotes, and GA represses dormancy.

### Reduced gibberellin levels promote secondary dormancy in Ae. arabicum

The transcriptional repression of genes involved in GA biosynthesis during light-induced dormancy and the lack of secondary dormancy in *rgl2* mutants, in which GA biosynthesis genes are derepressed, are consistent with the hypothesis that reduced GA activity is required for secondary dormancy. To test this hypothesis, we determined the impact of lowering GA levels by disrupting GA synthesis on secondary dormancy using paclobutrazol, a small molecule that inhibits GA synthesis ^22,23^. PAC-treatments were applied to seeds grown in 24 h dark for 7 days (conditions in which secondary-dormancy is not induced) and seeds were transferred to PAC-free plates in 24 h dark for 7 days and scored for germination (Fig. 4 B). PAC-treated seed did not germinate during the treatment and failed to germinate after the treatment on PAC-free plates, while controls treated with the solvent DMSO germinated (Fig. 4 A,B). This induction of secondary dormancy by PAC-inhibition of GA was evident after 24 hours of PAC-treatment (Fig. 4 C). The maintenance of dormancy after the removal of the PAC-treatment is consistent with the hypothesis that the inhibition of GA synthesis induced secondary dormancy (Fig. 4 B). Furthermore, PAC failed to induce dormancy in the *rgl2* mutant (Fig. 4 D), which indicates that the induction of dormancy by PAC required RGL2 function. If inhibition of GA synthesis promotes light-induced secondary dormancy, we predicted that light-induced secondary dormancy would be alleviated by GA treatment. As predicted, applying GA after the secondary dormancy was induced by 24 h light for 7 days alleviated dormancy (and promoted germination) (Fig. 4 E). Taken together, these data are consistent with the hypothesis that GA reduction by light induces secondary dormancy and GA-treatment alleviates light-induced dormancy.

**Figure 4.**
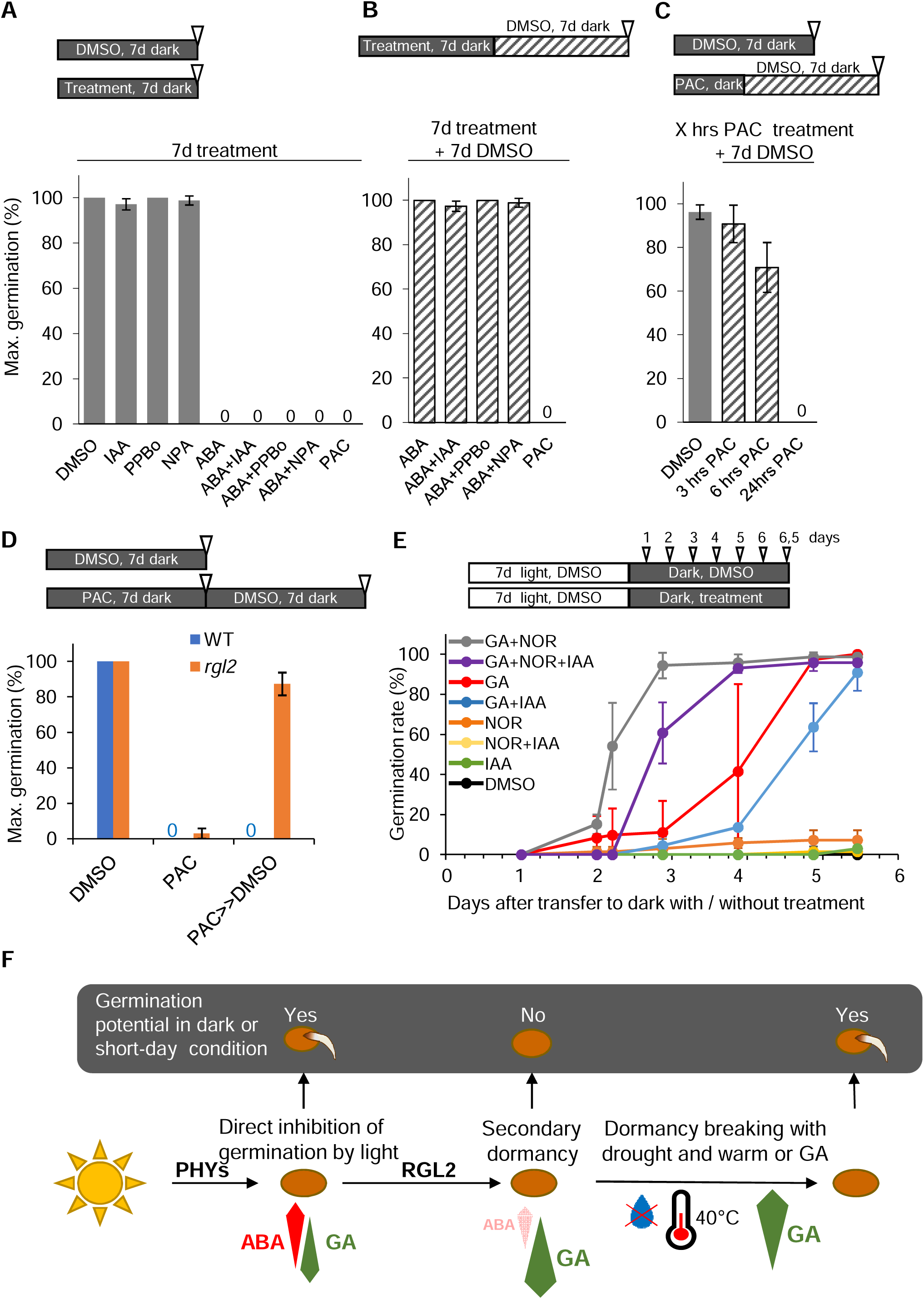
Hormone action in dormancy induction and alleviation in *Ae. arabicum* seeds. **(A)** Maximal germination scored after 7 days incubation of seeds in darkness on plates supplemented with IAA (auxin), PPBo (auxin synthesis inhibitor), NPA (auxin transport inhibitor), ABA (abscisic acid), PAC (gibberellin synthesis inhibitor) or DMSO as a control. **(B)** Potential dormancy induction by hormones and inhibitors was tested by scoring maximal germination of those samples not germinated in A after transfer to drug-free plates and 7 more days dark incubation. **(C)** Maximal germination after after 3, 6 or 24 hours of PAC treatment in darkness followed by 7 days on drug-free plates in darkness. **(D)** Maximal germination of WT and *rgl2* seeds after 7 days in darkness without or with 100 µM PAC, directly or after 7 more days without drug. **(E)** Dormancy alleviation after first establishing light-induced dormancy (7 days light as before), then transferring seeds to darkness on plates supplemented with either GA (GA_4+7_), NOR (ABA biosynthesis inhibitor), IAA (auxin), combinations, or DMSO as a control. Germination rate was scored over time; 0 days indicate the transfer. **(A-E)** Error bars: standard deviation of three biological replicates. **(F)** Hypothetical model of the light-induced secondary dormancy in *Ae. arabicum* seeds. Short light exposure (e.g., one day) inhibits the germination reversibly, allowing subsequent germination in darkness. Continuing light exposure gradually establishes secondary seed dormancy, mediated by RGL2 and induced by the downregulation of gibberellin biosynthesis. Secondary dormancy persists even after transfer to darkness, but can be alleviated to full germination capacity by the combination of heat and drought or by external addition of GA.

Since GA and ABA have opposite physiological effects on seed germination, we tested if ABA-treatments would induce secondary dormancy. Seed were treated with ABA in 24 h dark for 7 days before being transferred to media without ABA in the darkness (conditions that induce germination in controls). Treatment of seeds with ABA had no impact on secondary dormancy; seed germination on ABA-free plates was fully restored after the removal of the treatment (Fig. 4 B). Furthermore, norflurazon (NOR) – a carotenoid biosynthesis inhibitor used to inhibit ABA synthesis – did not revert secondary dormancy once it had been induced by exposure to white light for 7 days (Fig. 4 E). This indicates that reducing ABA activity had no impact on secondary dormancy. Since neither ABA-treatment nor reduction of ABA synthesis by NOR-treatment altered secondary dormancy, we conclude that ABA is not directly involved in establishing secondary dormancy. Similarly, treatment with auxin, auxin synthesis inhibitor (PPBo), or auxin transport inhibitor (NPA), alone or in combination with ABA, did not alter the secondary dormancy induction or alleviation (Fig. 4 A,B,E). Therefore, we found no evidence that ABA or auxin are involved in the establishment of secondary dormancy. Taken together, these data demonstrate that a light-induced reduction of GA synthesis – which is mediated in part by RGL2 – results in secondary dormancy.

## DISCUSSION

We demonstrate that secondary dormancy is induced by exposure of *Aethionema arabicum* seed to long day conditions and this may be an adaptation to Mediterranean climate. We show that light exposure of seed reduces GA synthesis which results in dormancy. The repression of GA required RGL2 activity; mutants that lack RGL2 function germinate after light exposure that induce secondary dormancy in wild type. Unexpectedly, ABA which plays a role in seed dormancy in many species, is not involved in the induction of secondary dormancy in *Ae. arabicum*. Importantly, light inhibits germination also in the *rgl2* mutant seeds as long as the seeds are kept in the light. This demonstrates that this direct light-inhibition of germination is mechanistically distinct from the long-lasting light effect that induces secondary dormancy; the transition from the direct light-inhibition to secondary dormancy depends on RGL2, likely repressing GA (Fig. 4 F).

The data are consistent with a mechanism for adaptation of *Ae. arabicum* to the Mediterranean climate characterized by hot, dry summers ^24,25^. Seed germination of *Ae. arabicum* is restricted to a short period in early spring, when the combination of short-day conditions, low temperature (11-14°C), and water availability are permissive for germination and seedling establishment (Fig. S1B) ^24^. We speculate that the induction of secondary dormancy during the early summer – when days are longer than in the early spring – blocks the germination of seeds which would otherwise face a hot and arid environment that is not amenable to seedling establishment. Subsequently, the exposure of secondary dormant seed to the heat and drought of the summer would alleviate secondary dormancy. This would ensure that germination of the seed in which secondary dormancy had been induced would be postponed to autumn or the following spring, when short days, permissive germination temperature and water accessibility again coincide. Failure of secondary dormancy induction would cause germination during summer, should permissive temperature and water availability occur for a short time due to stochastic weather conditions. The likeliness of following hot, dry days would result in high levels of seedling mortality. Annual dormancy cycling of seed populations is known to be regulated by temperature alone or in combination with water potential ^6,7,26–29^. Light is generally known to alleviate dormancy where darkness induces secondary dormancy in plants such as Arabidopsis ^2,8,29–31^. The response in *Ae. arabicum* operates opposite to Arabidopsis; light inhibits germination in *Ae. arabicum*, while darkness inhibits germination in Arabidopsis. Consequently, the secondary dormancy is induced by light in *Ae. arabicum* while it is induced by darkness in Arabidopsis. An *Ae. arabicum*-like mechanism may be general among species where light inhibits the germination, as demonstrated for *Brachypodium distachion* ^32^. We speculate that the mechanism reported here may operate in other species including crops and might have agricultural impact.

Our data demonstrate that it is the lack of GA and not an accumulation of ABA that promotes secondary dormancy. There is abundant evidence that GA promotes seed germination and represses primary dormancy in many species ^33,34^. Similarly, in *Ae. arabicum*, we demonstrate that inhibiting GA synthesis by PAC alone is sufficient to induce secondary dormancy. Furthermore, light-induced secondary dormancy can be alleviated by external GA treatment. Finally, secondary dormancy is defective in plants homozygous for a loss of function mutation in the *RGL2* gene, which encodes a repressor of GA responses. These data are consistent with the hypothesis that lack of GA promotes the development of secondary dormancy.

We have no evidence for a role of ABA in secondary dormancy in *Ae. arabicum*. ABA inhibits seed germination and is associated with seed dormancy in many species (Rodríguez-Gacio et al. 2009; Ali et al. 2021). Genes encoding enzymes of ABA synthesis, perception, and signaling have been shown to be part of a mechanism that promotes primary dormancy^35,37,38^, whereas negative regulators of ABA synthesis, such as protein phosphatase type 2C (PP2C) phosphatases, repress primary dormancy ^39,40^. Similarly, ABA promotes secondary seed dormancy in the *Cvi* of *Arabidopsis thaliana* accession ^4^. By contrast, ABA-treatment does not induce secondary dormancy in *Ae. arabicum* and light-induced secondary dormancy is not alleviated by inhibiting ABA synthesis. Furthermore, the steady state levels of mRNAs for ABA signaling genes *ABI3, ABI4, and ABI5* decrease during the development of secondary dormancy in *Ae. arabicum*. Therefore, the expression of genes involved in ABA signalling is inconsistent with ABA promoting secondary dormancy in *Ae. arabicum*. Repressing the synthesis of a germination promoter rather than promoting the synthesis of a molecule like ABA may be a plausible strategy for the induction of dormancy in seed, where metabolic activity will be low for an unpredictable period.

Our data demonstrate that light and long-day regime induce a robust and uniform secondary dormancy in *Ae. arabicum* (CYP accession) seeds. Secondary dormancy is a key mechanism for adaptive dormancy cycling in the seed bank of *Ae. arabicum*, and its establishment indicates that the responsible environmental triggers install a state that is maintained beyond the exposure to the trigger. Understanding the molecular control of secondary seed dormancy in wild plants provides a mechanistic explanation for a key event that influences life history dynamics in a changing environment. If such a molecular mechanism operates in seed crops, its manipulation has the potential to protect yields in the face of a changing global climate.

## MATERIALS AND METHODS

### Plant and seed materials

Experiments were conducted with *Aethionema arabicum* (L.) Andrz. ex DC. CYP (Cyprus) accession (obtained from Eric Schranz, Wageningen). The plants were propagated for seed material under 16 h light/19°C and 8 h dark/16°C diurnal cycles, under ∼300 μmol m^-2^ s^-1^ light intensity. WT and *rgl2* plants were randomly distributed on the shelves.

### Germination test

After seed harvest, seed stocks were kept in darkness at 50% humidity and 24°C for six months, except for the experiments in Fig. 1B, Fig. 2D and Fig. S1A where the seed age is indicated. One seed batch consists of the harvest from at least 6 plants; replicates represent different seed batches. Germination tests were conducted at the optimal temperature of 14°C in Petri dishes on 2-layer filter paper wetted with distilled H_2_O and supplemented with 0.1% Plant Preservative Mixture PPM (Plant Cell Technology) containing 0.135% 5-chloro-2-methyl-3(2H)-isothiazilone and 0.0412% 2-methyl-3(2H)-isothiazolone. Hormonal or chemical treatments were performed with GA_4+7_ (Duchefa), indole-3-acetic acid (IAA, Sigma-Aldrich), abscisic acid (Cayman Chemical, 10073), paclobutrazol (PAC, Duchefa), norflurazon (NOR, Sigma-Aldrich), 4-phenoxyphenylboronic acid (PPBo, Sigma-Aldrich), or naptalam (*N*-(1-Naphthyl)phthalamidic acid, NPA, Sigma-Aldrich). Unless indicated otherwise, 100 µmol IAA, PAC, NPA, PPBo, 10 µmol GA_4+7_, ABA, NOR, or 0.1 v/v % DMSO were applied.

### Light treatments

The respective light-dark regime and treatment scheme is indicated in all figures. Seed germination tests under white light were carried out in a Percival plant growth chamber equipped with fluorescent white light tubes with a spectrum from 400-720 nm (Figure S7) ^13^. Unless otherwise indicated, 130 μmol m^-2^ s^-1^ light intensity was used. For dark incubation, plates were transferred into a dark box covered with aluminum foil and black clothes and transferred back to the same Percival chamber. For diurnal regimes, the Percival growth chamber was equipped with a Valoya LightDNA-8 LED light source (https://www.valoya.com/lightdna/) and a clock timer. Light spectra and intensity were measured by LED meter MK350S (UPRtek). Red, far-red, and blue light treatments were performed using narrow-band LED light sources, peaking at 658 nm, 735 nm, and 450 nm, respectively (Fig. S7) ^13^. Licht intensity was 100 μmol m^-2^ s^-1^ for red and blue, and 10 μmol m^-2^ s^-1^ for far-red.

### Mutant screen and identification of the causative mutation in line p24H4

Generation and propagation of the mutant seed collection with 1320 lines were described in^13^. Germination in either M2 or M3 seed batches was screened for the loss of secondary dormancy after seven-day light exposure followed by seven-day darkness (7dL7dD). The p24H4 line originates from an M3 seed batch with 100% germination after 7dL7dD, indicating that the seed batch was homozygous for the causative mutation. After backcrossing one p24H4 plant with a WT CYP plant, the segregation of the F2 seed generation was tested for the secondary dormancy phenotype as described above (Fig. S2A). Twenty-eight seedlings germinated and represent the pool with the p24H4 mutant phenotype; the remaining 73 seeds were induced to germinate by GA treatment and represent the WT pool. DNA of both pools was isolated and sequenced on an Illumina H2500 platform with 100 bp single end mode by the Next Generation Sequencing Facility of the Vienna BioCenter Core Facilities (VBCF), member of the Vienna BioCenter (VBC), Austria. Sequencing reads were processed by the CLC Genomics Workbench 9.5.1 software (Qiagen). Reads were mapped to the *Aethionema arabicum* PacBio contigs originating from the CYP accession ^13^. After removal of duplicated reads, 122 and 125 million reads resulted in 72x and 74x coverage in the p24H4 and wild-type pool, respectively. Given the recessive inheritance of the mutant phenotype, the p24H4 pool was expected to be homozygous for the mutation and the wild-type pool to be heterozygous for the mutation or homozygous for the WT allele, summing up to an expected 33.3% mutant allele frequency. Variants were called with 90% minimum frequency in the p24H4 mutant pool and filtered out those which appear in the wild-type pool with higher than 40% frequency. 244 single nucleotide polymorphisms were found with these criteria and further filtered for overlap with coding regions of genes, leaving two mutations. One of them could be excluded due to mapping errors. The remaining mutation identified the gene *Aa31LG5G13950* that encodes RGL2. The mutation was confirmed by PCR with the primers 5’ GAGTCGAACGACACGAGACAC and 5’ CTACTTATAGCTCGAGCTACG and Sanger sequencing of the amplicon.

### RNA extraction and quantitative RT-PCR

Imbibed CYP WT or *rgl2* seeds were illuminated at 14°C with 130 µmol m^-2^ s^-1^ white light for 23 h (1dL) and seven days (7dL). Seeds with intact seed coats were collected for RNA extraction, with three biological replicates for each sample. RNA extraction, cDNA synthesis and quantitative qPCR were performed as described ^12^ using the primer pairs listed in Supplementary Table S2. The geometric mean of expression of Aethionema orthologues of POLYUBIQUITIN10 (*AearUBQ10, Aa3LG9G835*) and ANAPHASE-PROMOTING COMPLEX2 (*AearAPC2*, *Aa31LG10G13720*) was used for normalization ^12^. For each gene, the average expression levels of three replicates of 1dL WT seeds was set to one. Error bars represent standard deviation. Different letters on Fig. 3C indicate significant differences between any of the pairwise comparisons with p-value p<0.05 calculated with the Welch test.

### RNAseq analysis

Total RNA samples prepared as described above were used for RNAseq library preparation using Lexogen SENSE library kit with polyA selection. Libraries were sequenced with Illumina HiseqV4, single reads 50 mode. Acquired reads were processed with Trimmomatic to trim adapters ^41^. Reads were mapped against the *Aethionema arabicum* genome v3 (https://plantcode.cup.uni-freiburg.de/aetar_db/downloads.php) with default settings and `--no-unal --very-sensitive-local`. After removing duplicated reads, reads were counted with HTSeq-count with settings `-f bam -s no -t gene -i ID -m intersection-nonempty`, using the *A. arabicum* annotation v3.1 ^42^.

### GSEA analysis

Gene Set Enrichment Analysis (GSEA) was performed with clusterProfiler ^43^. Out of 24932 genes, an Arabidopsis orthologue was found for 20361 genes, and this was used to call GO terms ^42^. The gene set enrichment was calculated with “gseGO” function, using Benjamini-Hochberg adjusted *P*-value <0.05 and the log2 fold changes (LFC) of the RPKM values in four pairwise comparisons: WT 1dL vs. WT 7dL, *rgl2* 1dL vs. *rgl2* 7dL, WT 1dL vs. *rgl2* 1dL and WT 7dL vs. *rgl2* 7dL. Redundant GO terms were removed by the “simplify” function with default settings. The significant gene sets found in all four comparisons are listed in Supplementary Table S3.

### Data repository

RNAseq experiments, sequenced data files, and the processed data matrix have been deposited at the GEO as GSE237220 (<u>GEO Accession viewer (nih.gov)</u>. Processed data (RPKM values) of each sample and average values have been uploaded to the *Aethionema arabicum* Gene Expression Atlas (<u>Ae. arabicum Xpr (uni-freiburg.de)</u> as Dataset: 06 rgl2 mutant seeds light exposurè.

## Supporting information

Suppelemntal Table S1

## ACKNOWLEDGEMENT

We thank Eric M. Schranz for providing Aethionema seed stocks. We also thank the staff of the Vienna BioCenter Core Facilities GmbH (VBCF), a member of Vienna BioCenter (VBC), Austria, especially the Plant Sciences Facility for the growth of the plants, the Next Generation Sequencing Facility for generating the RNA sequencing data, the Molecular Biology Unit for providing multiple reagents, and the Vienna Covid-19 Detection Initiative (VCDI) for generating a safe work environment during the pandemic. We thank Nicole Lettner for the technical support.

## Author Contribution Statement

Z.M., K.G., G.L.-M., O.M.S. and L.D. conceived the project and conceptualized the work. Z.M., F.X. and K.G. performed the experiments; Z.M., K.G. and F.X. prepared and handled samples; Z.M., P.K.I.W., S.A.R., N.F.P., M.D. and K.L. performed the transcriptomics data analysis. Z.M., O.M.S. and L.D. wrote the manuscript. All authors read and commented on the manuscript.

## SUPPLEMENTARY MATERIAL

**Supplementary Figure S1.**
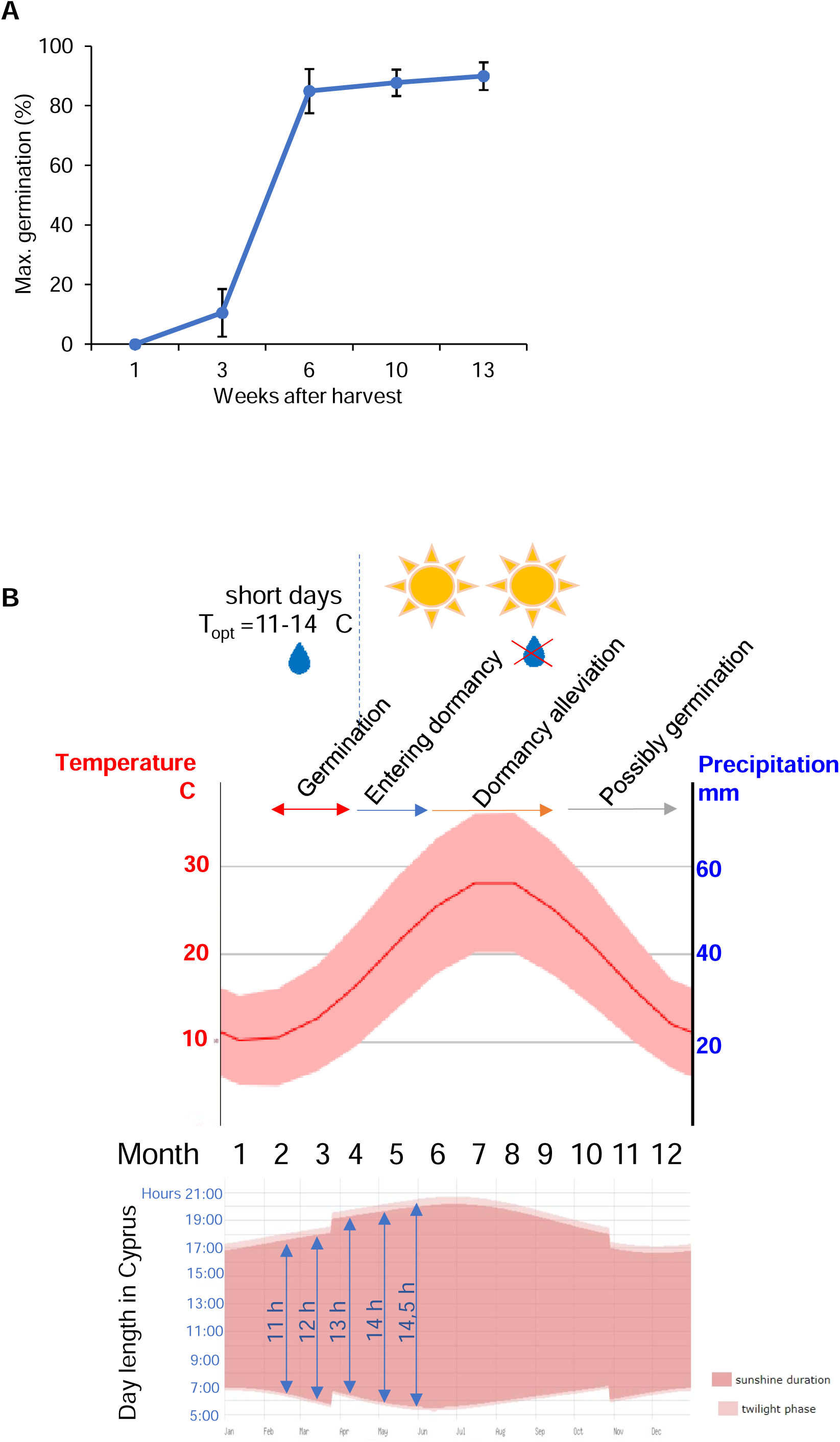
(A) Release of primary seed dormancy in wild type seeds (B) Annual dormancy cycling of *Aethionema arabicum* (CYP) corresponding to the climate data. Source of climate data: https://en.climate-data.org/.

**Supplementary Figure S2.**
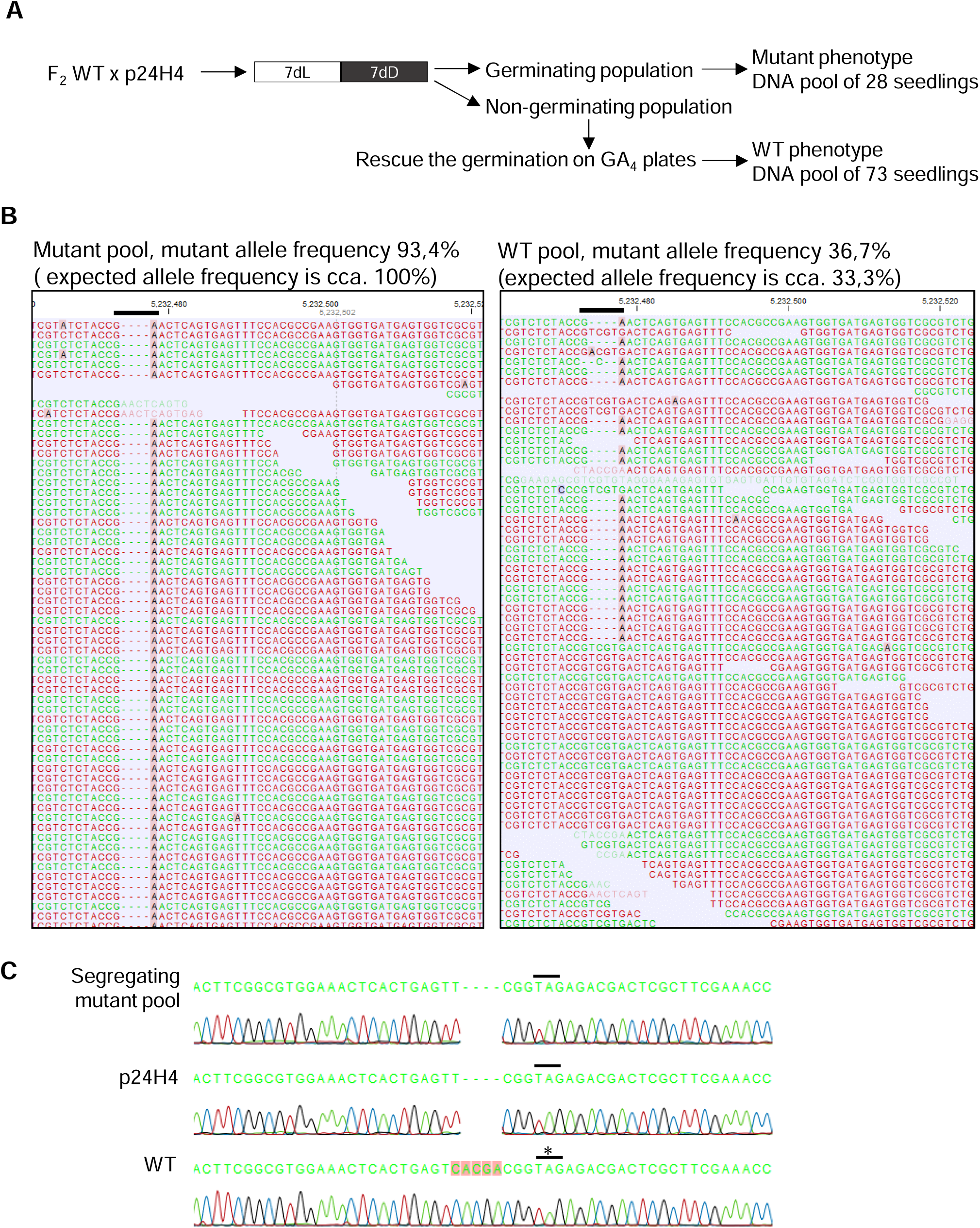
Identification of the mutation in p24H4 by whole-genome sequencing of segregating population.

**Supplementary Figure S3.**
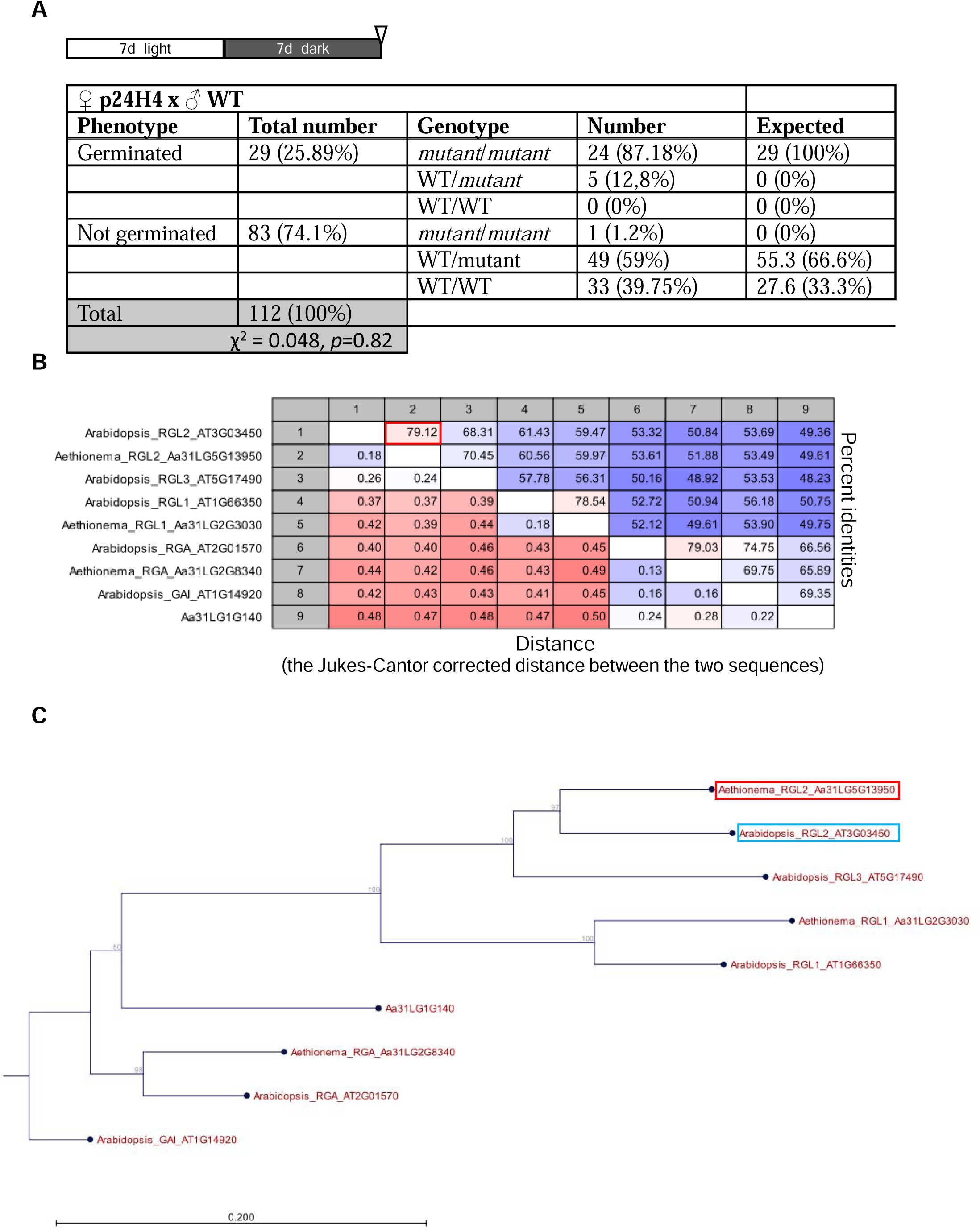
Confirmation of the *rgl2* mutant (A) Co-segregation analysis of the mutant secondary dormancy phenotype with the identified mutation in the *RGL2* gene. (B) Pairwise comparison of the DELLA proteins. (C) Phylogenetic tree of DELLAs in *Ae. arabicum* and Arabidopsis.

**Supplementary Figure S4.**
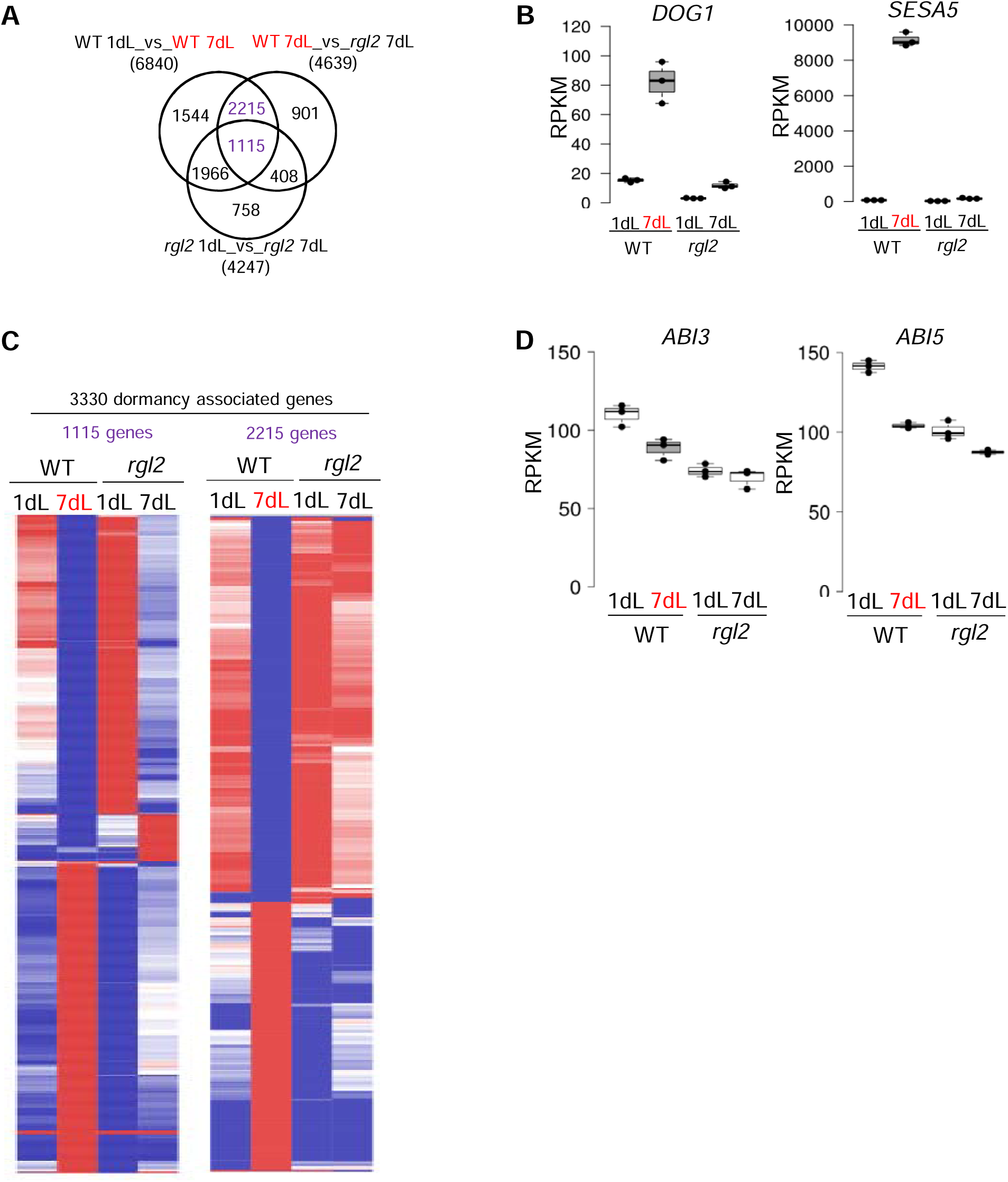
Transcriptome identifies secondary dormancy associated genes.

**Supplementary Figure S5.**
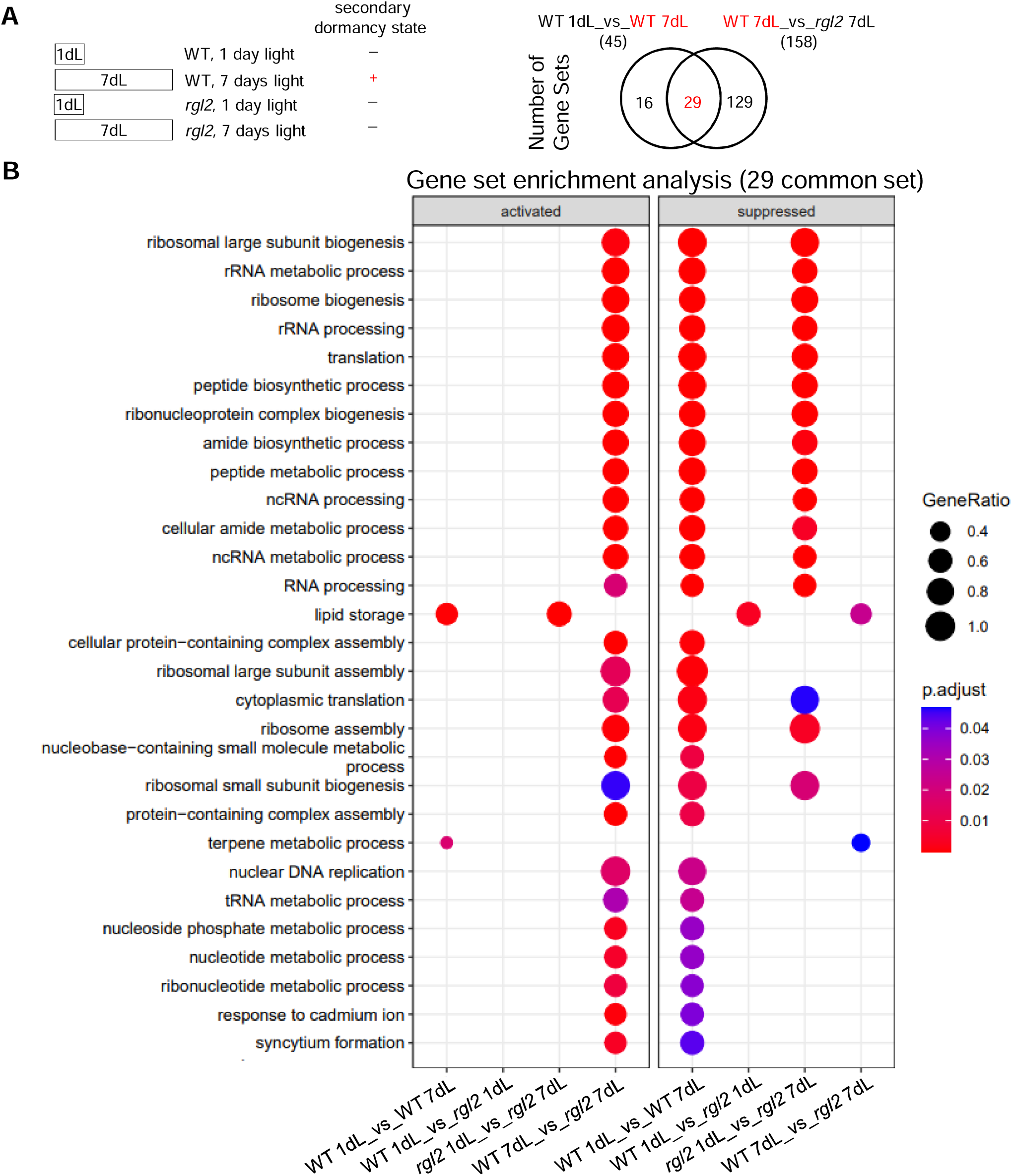
Gene set enrichment analysis of the light induced secondary seed dormancy in wild type versus *rgl2* mutant.

**Supplementary Figure S6.**
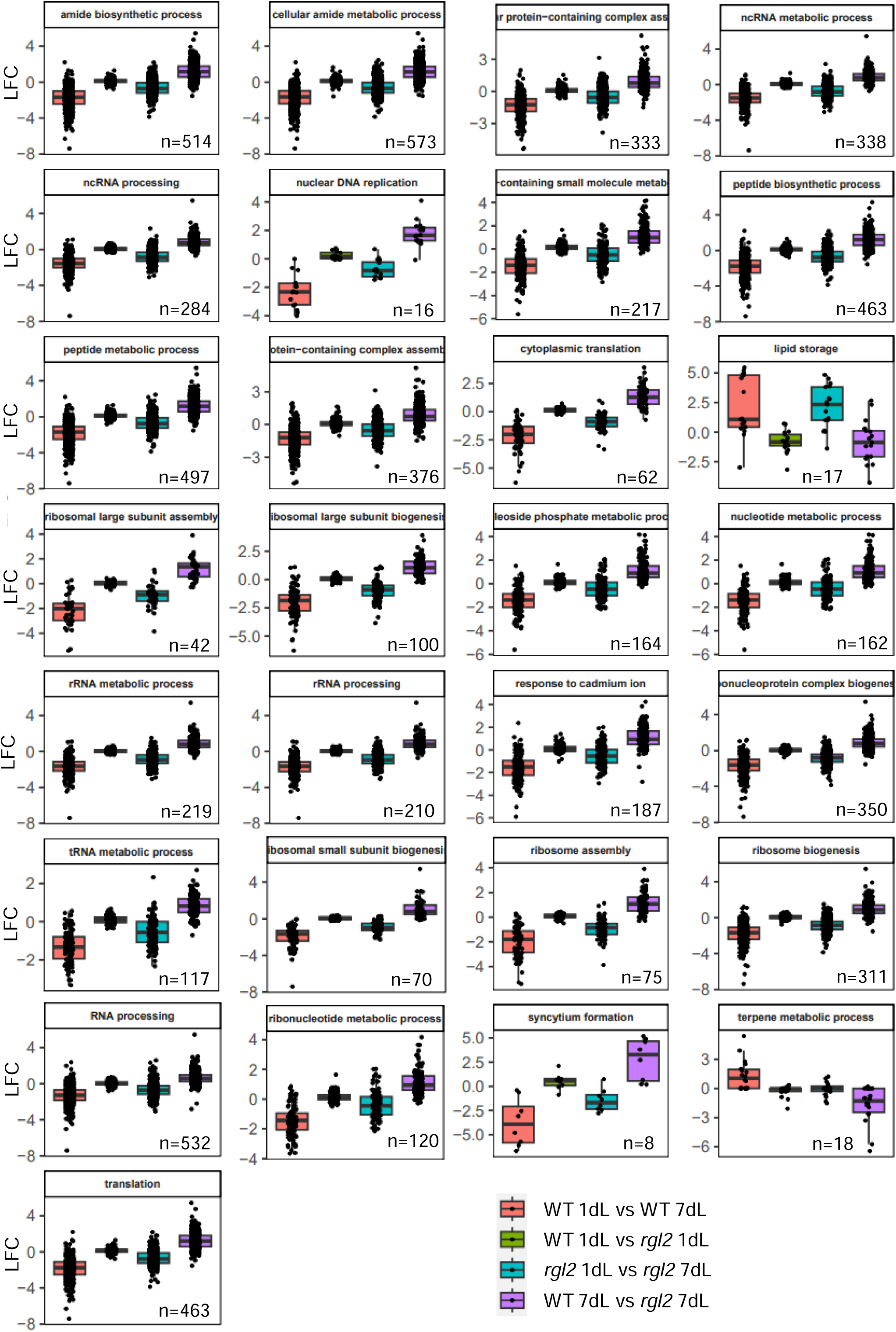
Expressional changes of the 29 gene sets common in non-dormant versus dormant comparisons.

**Supplementary Figure S7.**
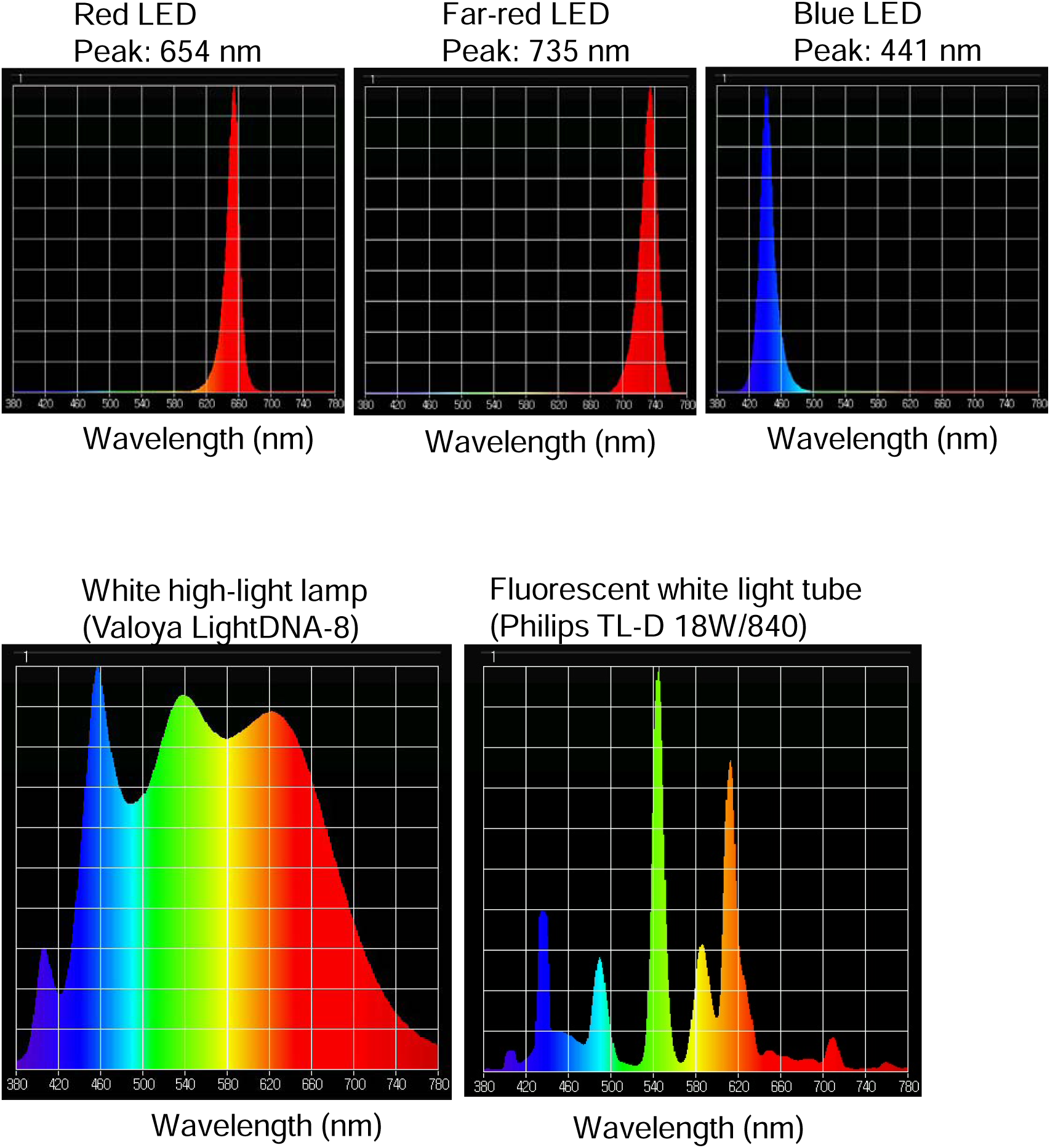
Spectral properties of the used light sources.

**Supplementary Table S1.** List of all DEGs identified in four comparisons and the list of 3330 genes identified as dormancy associated genes.

**Supplementary Table S2.** List of primers and accession number of *Aethionema arabicum* genes used for quantitative RT-PCR.

**Supplementary Table S3.** List of all gene sets identified in four comparisons.

**Supplementary Table S4.** List of the log fold changes of all gene expression in the 29 common gene sets.

