## Supplemental Table S2 for "Long days induce adaptive secondary dormancy in seed of the Mediterranean plant *Aethionema arabicum*"

**Supplemental Table S2 List of primers used for quantitative RT-PCR analysis**

| **Name** | **Nucleotide sequence** | **Accession number v3.1** |
| --- | --- | --- |
| AearUBQ10_for | GAGGATGGCCGAACATTG | *Aa3LG9G835 (v3.0)* |
| AearUBQ10_rev | TGCCCGTTAGGGTTTTGA |  |
| AearAPC2_for | TCTCCTGCAATCGAGGACTT | *Aa31LG10G13720* |
| AearAPC2_rev | GCAGTGAGCAACCGGTATTT |  |
| AearDOG1_for | CGCGTCACTAAGCGATCTAAC | *Aa31LG9G7230* |
| AearDOG1_rev | GCCGCGTCTTCTTGTAGACTT |  |
| AearSESA5_for | TGCTCTCTTCCTCCTCCTTG | *Aa31LG3G10660* |
| AearSESA5_rev | CATTTGCCTTGTTGTTGTGG |  |
| AearGA3ox1_for | TCTTCGTCACCTCCCTGACT | *Aa31LG7G270* |
| AearGA3ox1_rev | GATGAGCGGGAGAGTTGTGT |  |
| AearNCED6_for | GCTTCTTCAGCTCTCGACAA | *Aa31LG8G10550* |
| AearNCED6_rev | GAACCGTTGGATCAGTCGGT |  |
| AearRGL2_for | GGACCCTGCAACAATACCAT | *Aa31LG5G13950* |
| AearRGL2_rev | CCACGCCTTCAACTTCCTTA |  |
